# Soybean aphids exploit abscisic acid signaling to suppress jasmonate defense responses

**DOI:** 10.1101/2025.09.21.677601

**Authors:** Jessica D. Hohenstein, Charles Kanobe, Martha I. Natukunda, Patricia Gallardo, Dandan Zhang, Nik Kovinich, Anjel M. Helms, John F. Tooker, Gustavo C. MacIntosh

## Abstract

**Summary:**

- Soybean aphids (*Aphis glycines*) can induce susceptibility on soybean (*Glycine max*) during colonization. However, the mechanism for this process is not known. Based on previous transcriptome analyses, we hypothesized that aphids block effective jasmonate (JA) defenses by inducing an antagonistic abscisic acid (ABA) signal.
- To test this hypothesis, we used a combination of gene expression analyses, measurements of hormone levels, and aphid bioassays on plants with reduced expression of ABA-related genes.
- Aphid feeding attenuated JA responses in soybean plants and facilitated the growth of a chewing herbivore. Aphid-treated plants had increased levels of *cis*-JA but not biologically active JA-isoleucine, and aphid feeding induced expression of genes associated with JA-Ile catabolism. In parallel, aphid-feeding induced higher levels of ABA. ABA treatment and knockdown lines impaired in ABA biosynthesis (*aba2*-RNAi) or signaling (*scof-1*-RNAi), showed that ABA suppressed wound-induced JA responses. Aphid populations were significantly reduced on ABA-deficient plants and aphid-regulated attenuation of JA signaling was abolished in these lines. Remarkably, plants defective in ABA signaling had increased JA signaling in the absence of stressors.
- Our results indicate that, in soybean, the ABA pathway is necessary to control basal levels of JA and soybean aphids exploit this ABA-JA antagonism to suppress plant defenses.

## Introduction

Unlike plant-chewing insects that cause significant mechanical damage to plants, phloem-feeding insects such as aphids minimize tissue damage using piercing-sucking mouthparts (i.e., stylets; (Walling 2008). Perception of aphids by plants is thought to include detection of salivary components, chitin fragments from the exoskeleton, or physiological changes in sieve elements (Libault *et al*. 2007; Hohenstein *et al*. 2019; Kaloshian and Walling 2016; Will and van Bel 2006; Hogenhout and Bos 2011). Phytohormones play a significant role regulating plant responses to hemipteran infestations, but effects of individual hormones seem different for specific plant-hemipteran interactions, and highly context-dependent (Erb and Reymond 2019; Åhman *et al*. 2019). Jasmonic acid (JA), however, is accepted as a positive regulator of plant defenses against phloem-feeding insects. Increased plant resistance after treatments with JA or MeJA has been observed in several systems, including increased resistance to green peach aphid in Arabidopsis (Ellis *et al*. 2002), to potato aphid in tomato (Cooper and Goggin 2005), to greenbug aphid in sorghum (Zhu-Salzman *et al*. 2004), to mustard aphid in Indian mustard (Koramutla *et al*. 2014), and to grain aphid in wheat (Cao *et al*. 2014). Genetic approaches have resulted in similar conclusions, as mutants defective in JA signaling are usually more susceptible to aphid infestation while plants overexpressing JA biosynthesis or signaling genes are more resistant (Losvik *et al*. 2017; Mewis *et al*. 2005; Mewis *et al*. 2006; Ellis *et al*. 2002; Kersch-Becker and Thaler 2019).

Phloem feeders can in return suppress effectual defenses including those regulated by JA (Kaloshian and Walling 2016; Hogenhout and Bos 2011; Åhman *et al*. 2019). For example, pea aphid infestation suppressed JA levels and accelerated development of aphid nymphs on broad bean plants (Takemoto *et al*. 2013). Moreover, cabbage aphid infestations suppressed JA accumulation and signaling in wild cabbage, facilitating growth and development of *Pieris brassicae* caterpillars after sequential colonization (Soler *et al*. 2012). Experiments analyzing responses to co-infestation with aphids and caterpillars in two closely related milkweeds showed that oleander aphids can attenuate caterpillar-induced JA responses in *Asclepias syriaca* but not in *A. tuberosa*, indicating that suppression of defenses is determined by factors in both aphids and host plants (Ali and Agrawal 2014).

Hemipterans may exploit antagonism among phytohormone signaling by eliciting a decoy response. Induction of the SA pathway by silverleaf whitefly feeding can suppress JA defenses in Arabidopsis, facilitating their own establishment (Zarate *et al*. 2007). Similarly, by manipulating SA-JA crosstalk, solenopsis mealybug suppressed JA defenses, including production of gossypol, which has insecticidal activity (Zhang *et al*. 2011; Zhang *et al*. 2015; Ma *et al*. 2019). Green peach aphids, on the other hand, induced accumulation of ABA and aphid performance was reduced on ABA-deficient mutants, suggesting that they can manipulate effective defenses by inducing ABA signaling that antagonizes production of indole glucosinolates (Hillwig *et al*. 2016). However, not all ABA mutants of Arabidopsis had increased susceptibility to aphids (Hillwig *et al*. 2016; Kerchev *et al*. 2013), suggesting tight regulation of different branches of ABA pathways. Lastly, hemlock woolly adelgids may suppress JA-induced defenses in Eastern hemlock by exploiting both SA-JA and ABA-JA antagonism (Schaeffer *et al*. (2018).

Soybean aphid (*Aphis glycines* Matsumura) is an invasive specialist phloem-feeder pest first reported in the US in 2000. Rapid reproduction, withdrawal of plant nutrients, and their ability to vector plant viruses make this insect species an important pest of soybeans in the US Midwest (Tilmon *et al*. 2011). Resistant soybean cultivars (*Rag*) are available, and currently 12 loci associated with host-plant resistance to soybean aphid have been identified, although the genes conferring this resistance have not been isolated (Natukunda and MacIntosh 2020). Transcriptome and metabolite analyses have shown that, upon detecting aphids via recognition of herbivore-associated molecular patterns (HAMP), perhaps through the detection of chitin (Kaloshian and Walling 2016; Hohenstein *et al*. 2019), susceptible soybean plants activate basal defenses that include production of deterrent isoflavones and callose deposition (Hohenstein *et al*. 2019; Yao *et al*. 2020). Both JA and SA seem to have a positive role in regulating soybean defenses against soybean aphid (Selig *et al*. 2016; Yates-Stewart *et al*. 2020; Studham and MacIntosh 2013).

Soybean aphids can also manipulate plant defenses to increase host susceptibility (Varenhorst *et al*. 2015b; Varenhorst *et al*. 2015a). However, the molecular mechanisms mediating this effect are not known. Our previous transcriptome analysis showed that virulent aphids initially (1 day post-infestation) triggered induction of JA synthesis and the downstream response to the hormone. However, at longer colonization periods (7 days post-infestation), while transcripts associated with JA synthesis were still highly induced, transcripts associated with JA responses had similar expression levels in infested and control plants. These results suggested an aphid-triggered block of the ability of soybean plants to respond to increased JA levels (Studham and MacIntosh 2013). We also observed large increases in expression of ABA biosynthesis and ABA responsive genes. We hypothesized that aphids exploit ABA-JA antagonism via defense signaling to increase their feeding and colonization success (Studham and MacIntosh 2013).

Here, we tested this hypothesis. We demonstrate that aphids attenuate the ability of plants to respond to JA-mediated stimuli and that their reduced response has implications in plant-mediated pest interactions. We further show that knockdown of genes involved in ABA biosynthesis (*ABA2*) and signaling (*SCOF-1*) significantly reduced aphid populations. In unstressed, aphid free plants, ABA-deficiency also increased basal levels of JA signaling, indicating that aphids exploited the endogenous ABA-mediated regulation of basal JA response. We thus provide genetic evidence that aphid-mediated attenuation of JA defense responses is mediated by the endogenous ABA pathway in soybean. These findings provide new insights into aphid suppression of plant defenses through manipulation of defense signaling, highlighting potential targets for improving plant resistance and controlling aphid pests.

## Materials and Methods

For all experiments, aphid-susceptible soybean (*Glycine max* (L.) Merr.) seeds cv. SD01-76R were sterilized overnight using chlorine gas as previously described (Paz *et al*. 2006). Plants were grown in a growth chamber in steam sterilized Metro-Mix® 900 soil (Sun Gro Horticulture, Vancouver, BC, Canada) at a constant temperature of 25°C with a 16 light:8 dark photoperiod. Plants were fertilized once per week with 1 L of a 1:1 mixture of 6% All-Purpose Scott’s Miracle-Gro Excel (21-5-20, The Scott’s Company LLC, Marysville, Ohio, USA) and 6% Cal-Mag Miracle-Gro Professional (15-5-15, The Scott’s Co.) applied at a rate of 12.5 mL L^-1^ water.

### Aphid infestations

Soybean aphids (*Aphis glycines* Matsumura) virulent biotype 1 were obtained from a laboratory colony at Iowa State University. For aphid population quantification experiments, ten 5-6d old age-synchronized aphids were placed on one leaflet of the V3 or V4 trifoliate using a fine tip paintbrush and were confined using a clip cage (BioQuip products, Rancho Dominguez, CA, USA). For caterpillar, wounding and JA experiments that required “aphid pre-treatment”, thirty 5-6 d old age-synchronized aphids were used to colonize plants. For all experiment sets, plants without aphids also had clip cages attached to the leaf to account for any cage effect on the plant. Aphids were allowed to feed and reproduce for 7 days and then populations were counted. Both aphid-treated and control leaflets were gently brushed to remove aphids or mimic any mechanical stimulation caused by brushing.

### Caterpillar, wounding and JA treatments

Ten control or aphid pre-treated plants were used for each treatment. The three experimental conditions for the caterpillar experiments included: i) control (no aphids, no caterpillars), ii) caterpillar, and iii) aphid+caterpillar. Clip cages were placed on each infested leaflet to confine aphids and caterpillars to specific leaves. <u>For the</u> aphid+caterpillar treatment, aphids were allowed to feed on plants for 7 days, then aphids were removed using a fine paintbrush. One second instar larva corn earworm (*Helicoverpa zea*) caterpillar was placed on the same leaf. The weight of each caterpillar in grams was recorded before placement on the leaf and after feeding for three days. The normalized total weight gain was calculated for each caterpillar treatment group. At the end of the 3-day caterpillar treatment, leaf tissue was collected for the control, caterpillar, and aphid+caterpillar treatments for gene expression analysis. In separate experiments, plants were wounded with tweezers and controls were left untreated, or were sprayed with a solution of JA (1.5mM) dispersed in water from a stock of 1 g of JA per ml of acetone (Thaler *et al*. 1999), using a SureShot Atomizer Sprayer (Milwaukee Sprayer, Menomonee Falls, WI, USA) while control plants were treated with a similar solution of water:acetone without JA. Six hours after each treatment, leaves were collected for gene expression analysis.

### Hydroponic ABA treatment experiments

For hydroponic experiments, plants at unifoliate stage were gently removed from loose soil. Roots were rinsed in water and plants were placed on platforms with the roots submerged in modified Hoagland’s hydroponic media containing 1.25 mM KNO_3_, 2.16 mM Ca(NO_3_)_2_·4H_2_O, 0.75 mM MgSO_4_·7H_2_O, 0.3 mM KH_2_PO_4_, 50 µM KCl, 50 µM H_3_BO_3_, 10 µM MnSO_4_·H_2_O, 2 µM ZnSO_4_·7H_2_O, 2.4µM CuSO_4_·5H_2_O, 100 µM EDTA-Na_2_, 100 µM FeSO_4_·7H_2_O. Three plants were included in each container. Media was fully replaced once weekly and the volume of media was maintained by adding deionized water as needed. Plants were allowed to grow until the V3 stage (Fehr and Caviness 1977) before treatments were administered. Four treatments were administered: 0µM ABA, 0µM ABA+wounded, 100µM ABA, 100µM ABA+wounded. Each treatment had 5 replicates for a total of 20 containers, with 3 plants each. A 100mM ABA stock was dissolved in 100% methanol. When plants reached the V3 stage, hydroponic media was replaced with media supplemented with control (0µM, 0.1% methanol) solution or 100 µM ± ABA-supplemented solution (Sigma-Aldrich, St. Louis, MO). After 24h hours, V1 leaves were wounded with a tweezers or were left unwounded (control plants) within each ABA level. Six hours after wounding, V1 leaves of three plants in the same container were pooled for a total of 5 replicates per treatment. Samples were immediately frozen with liquid nitrogen and stored at-80°C until further sample processing.

### Vector construction and viral inoculation

Bean Pod Mottle Virus (BPMV) RNA components as well as Soybean Mosaic Virus (SMV) helper component were a kind gift from Dr. Steven Whitham (Iowa State University, Ames, IA) and Dr. Michelle Graham (USDA-ARS CICGR, Ames, IA). To generate *aba2* RNAi and *scof-1* RNAi constructs, approximately 300bp fragments of *ABA2* and *SCOF-1* were amplified by qRT-PCR (primers listed in Supporting Table S1) and then digested with *BamHI* and *XhoI* and ligated into the RNA2 vector. Construct sequences were confirmed and plants were bombarded, grown, tested for BPMV presence, stock tissue collected and stored according to Whitham *et al*. (2016). A slurry was made using 30mg stock tissue per 2mL of 50mM phosphate buffer pH 7.0. Carborundum (320-grit) was sprinkled onto the unifoliate leaves of 7-or 8-day old soybean seedlings. Then, 15µL of the slurry supernatant was pipetted onto each unifoliate leaf and rubbed gently taking care not to rip large holes in leaves. Plants were allowed to grow for approximately 3 weeks to allow for viral spread and gene silencing.

### Virus-induced gene silencing experiments

Ten to fifteen SD01-76R seeds were planted in one pot (Poly-tainer^TM^ #2, Nursery Supplies Inc., Orange, CA). After one week, seedlings were thinned to 4 per pot. At the unifoliate stage, plants were dark-treated for approximately 24 hours and the chamber temperature was changed to 21°C/18°C to facilitate viral infection and spreading.

### Experimental design, wounding and aphid treatments, and tissue collection for VIGS experiments

A Randomized Complete Block Design (RCBD) was used. For aphid phenotyping experiments, each construct was represented once in each pot with a total of 4 constructs (mock, virus vector control, *aba2* RNAi, and *scof-1* RNAi). Each experiment consisted of 13-15 biological replicates per construct or BPMV treatment. For wounding experiments, each pot contained four plants of the same construct. Each plant in the pot received a different treatment: control (C), aphid-infested for 7 days (A), wounded for 6 hours (W), or aphid-infested for 7 days and wounded for 6 hours (AW). Each experiment consisted of 6 biological replicates per construct and samples were collected by pooling two leaflets of the same treatment to generate 3 biological replicates for gene expression analysis. Aphid population quantification and wounding experiments were performed as described above. From the aphid population quantification experiments, leaf tissue from 2-4 plants with or without aphids was pooled and collected into liquid nitrogen for hormone analysis. All samples were stored at-80°C until further processing.

### Sample processing and gene expression quantification

All samples were ground in liquid nitrogen using a mortar and pestle. Total RNA was extracted from leaves using TriReagent (Ambion®, Life Technologies, NY, USA). Genomic DNA contamination was removed using TURBO DNA-*free*^TM^ kit (Ambion®, Life Technoligeis, NY, USA) and PCR/gel electrophoresis was conducted to check for any undigested contaminating genomic DNA. One microgram of cDNA was synthesized using qScript™ Flex cDNA Synthesis Kit (Quanta Biosciences, Beverly, MA, USA) using Oligo dT primers. Quantitative PCR (qRT-PCR) was done using PerfeCTa® SYBR® Green FastMix®, Low ROX (Quanta Biosciences Beverly, MA, USA) in an Mx4000 (Stratagene, Agilent, Technologies, Santa Clara, CA, USA). Cycle threshold values were quantified and analyzed according to a standard curve and then normalized to internal control gene *Glyma20g27950* (*UBQ*). All primers used in this study can be found in Supporting Table S1.

### Phytohormone quantification

ABA was extracted and quantified using a method adapted from (Forcat *et al*. 2008). Standards and anhydrous acetic acid were purchased from Sigma-Aldrich (Sigma-Aldrich, St. Louis, MO). Solvents were LC-MS grade (Fisher). Briefly, lyophilized soybean tissues were pulverized with 5 mm stainless steel grinding beads in a Mixer Mill MM400 (Retsch) equipped with an Adapterrack PTFE pre-frozen at-20°C. The fine powder (5 mg) was extracted on ice for 30 min in 400 µl of methanol:acetic acid:water (10:1:89) that contained 5µM (+)-catechin as an internal standard because of its ionization potential and absence from soybean (Kovinich *et al*. 2011). The supernatant was removed by centrifugation and the extraction repeated once as indicated above. The pooled supernatants were flash frozen in liquid nitrogen and lyophilized to dryness. The residue was resuspended in 100 µl of methanol:acetic acid:water (10:1:89), filtered by centrifugation through 2 µm PTFE for analysis by LC-MS.

For direct infusion mass experiments, a Q-Exactive Orbitrap mass spectrometer (Thermo Scientific, San Jose, CA, USA) was used to obtain exact mass measurements and parameter optimization prior to UHPLC-MS analysis. Hormone standards were prepared at 20 uM in 1:1 water:acetontirle. Standards were introduced individually by direct infusion through a heated electrospray source inlet (HESI) using a 4.00 kV spray voltage bias relative to the entrance orifice of the mass spectrometer. The capillary and Aux gas temperatures were programmed at 250°C and 50°C, respectively. The generated ions passed through the S-lens ion guide (60.0 V) and were subsequently transferred into an Orbitrap mass analyzer. The Orbitrap was scanned from m/z 80.0 to 600.0 with a resolving power of 70,000. A target AGC of 1.0×10^6^ with a 200 ms injection time was used for the analysis. These parameters were found to give maximum signal across all standard infusion experiments. The exact *m*/*z* values of each standard were recorded and used to generate an inclusion list for LC-MS experiments (see below).

For liquid chromatography-mass spectrometry analyses, a Q-Exactive Orbitrap mass spectrometer interfaced to an Accela UHPLC (Thermo Fisher Scientific, San Jose, CA) was used. Standards were analyzed in order to record retention times and to generate calibration curves prior to sample analysis. Standards were injected onto the system via the PAL autosampler (5µL) at concentrations 5, 2.5, 1, 0.5, and 0.1 µM. The UPLC was programmed for gradient delivery of water (solvent A) and acetonitrile (solvent B) each containing 0.1% acetic acid at 300 µL min^-1^. The gradient for solvent B was from 10-25% 0-2 min, 25-35% 2-7 min, 35-100% 7-9 min, 100% 9-11 min, 10% 11-12 min, held at 10% until 15 min. Separations were achieved using an Acquity UPLC BEH Shield RP18 analytical column (Waters) with a pore size of1.7 µm. The column held at 35°C using a column heater (Thermo Fisher Scientific, San Jose, CA). Analytes separated by the column were directed to the HESI source using the parameters discussed above. The Q-Exactive was operated in Selected Ion Monitoring mode, via the inclusion list generated from direct infusion experiments (see above). That is, the mass analyzer was programmed to scan each elution window and set to record ion intensity for the defined standard *m*/*z* values. For sample analysis, injection volumes were normalized to the amount of internal standard using Xcalibur software and then to the amount of dry tissue weight using Excel. ABA extraction and quantification was also confirmed using the protocol described in Schmelz *et al*. (2004).

### Statistical Analysis

Experiments were set up using either completely randomized design where t-tests (t-test assuming unequal variance) were used to determine significance or a randomized complete block design (RCBD) was used and data was analyzed by ANOVA analysis, as indicated in each figure. If an overall significant difference was detected, pair-wise comparisons were made using least significant difference (LSD) multiple comparison correction. All RCBD statistical analyses were done using Statistix9 software.

## Results

### Soybean aphids attenuate JA-, wound-, and herbivore-induced JA signaling and facilitate chewing herbivore performance

To test whether soybean aphids can attenuate JA responses in susceptible soybeans, we analyzed expression of *PinN2*, a JA-inducible marker gene (Botella *et al*. 1996) in control and aphid-infested plants following mechanical wounding or feeding by *Helicoverpa zea* caterpillars. For the aphid treatment, plants were infested with aphids for 7 days.

Aphid infestation alone had a small effect on induction of *PinN2* (Fig. 1a). Mechanical wounding (Fig. 1a) and caterpillar feeding (Fig. 1b) highly induced expression of *PinN2* in aphid-free plants. However, in aphid-colonized plants, induction of *PinN2* expression after wounding or caterpillar feeding was significantly attenuated when compared with aphid-free plants (Fig. 1a,b). Furthermore, *H. zea* caterpillars gained more weight after feeding on plants that had previously been infested by aphids (Fig. 1c). These results indicate that aphids blocked the accumulation and/or the perception of JA.

**Figure 1.**
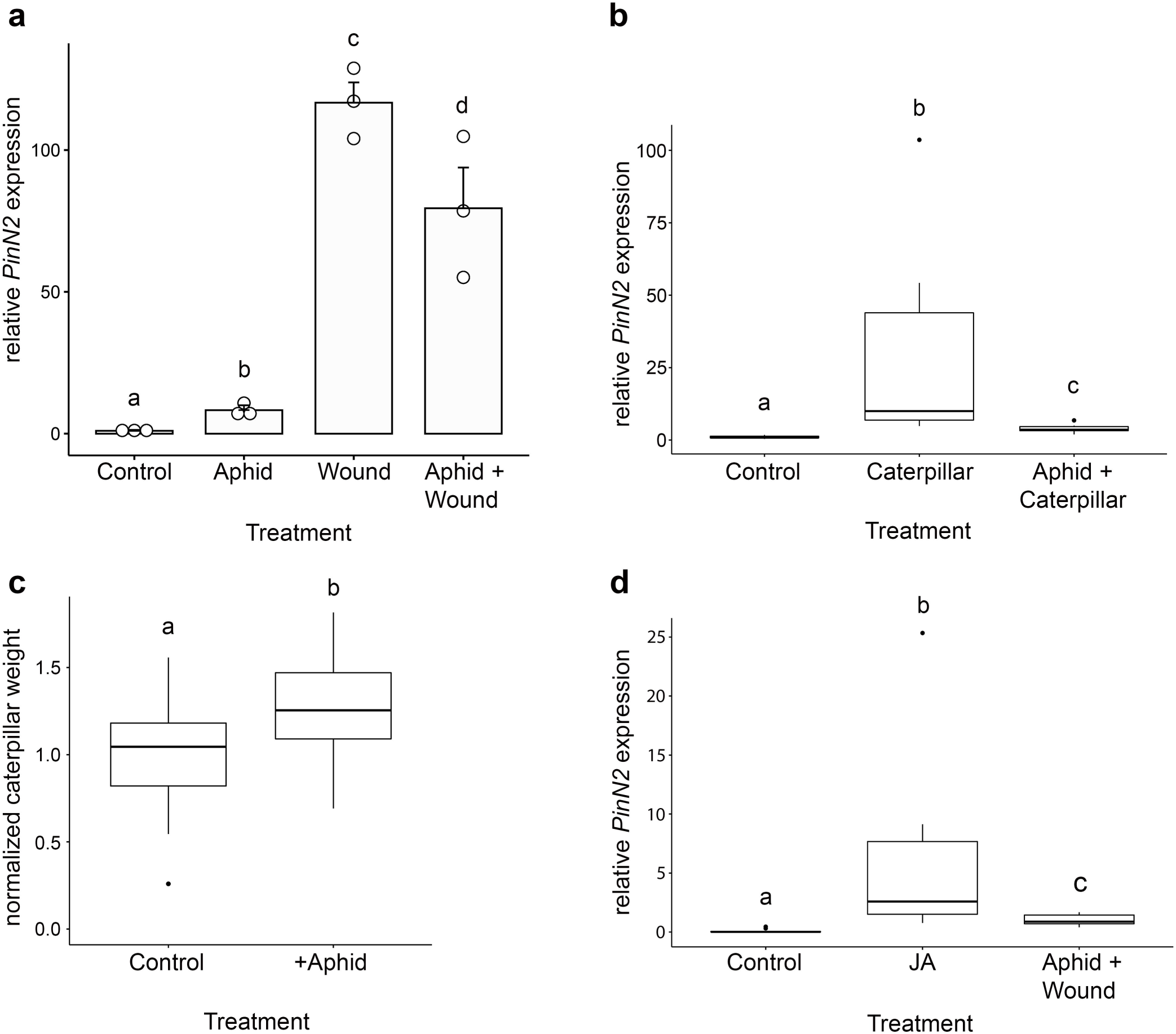
Soybean aphids suppress the plant JA response and facilitate a chewing herbivore. Plants were exposed to aphids for 7 days (Aphid) or were left unexposed (Control). Plants were wounded with tweezers (**a**), treated with JA (**b**) or exposed to caterpillar feeding (**d**). Six hours after treatment, samples were collected. Relative expression of *PinN2* was assayed using quantitative PCR. Growth of caterpillars, as weight gain, was measured on aphid-infested and control plants 3 days after applying caterpillars (**c**). Different letters indicate significance at P<lt;0.05 according to two-tailed unpaired t-test. Error bars in A represent standard error, mean for 3 independent experiments are indicated by circles. In box plots, the line inside the box represents the median value. The box represents the first to third quartiles. Bars indicate data distribution, and dots represent outliers. For box plots, results from at least 3 independent experiments were included.

The aphid-regulated attenuation of JA signaling could be due to reduced production of JA. Previous analyses in soybean showed that aphid infestation caused reduced levels of linolenic acid, the precursor to JA (Kanobe *et al*. 2015). To test if aphids regulate JA defenses via the reduction of JA levels, we directly applied JA to control and aphid-infested plants and measured *PinN2* marker gene expression. We hypothesized that if aphids block JA synthesis, the exogenous application of JA to plants would override the aphid’s ability to attenuate the response to the hormone. However, JA application resulted in a similar attenuation of *PinN2* induction by aphid feeding (Fig. 1d) compared to wounding (Fig. 1a). Aphid-infested plants treated with JA still had significantly lower *PinN2* expression when compared to JA-sprayed, aphid-free plants. Thus, the attenuation of *PinN2* induction after wounding in aphid-infested plants is due, at least in part, to a block in JA signaling downstream of JA biosynthesis.

### JA, but not JA-Ile accumulates in aphid-treated plants

To investigate potential steps in JA signaling that could be affected by aphid colonization, we first measured changes in levels of different jasmonates in response to aphid feeding. Aphids did not alter levels of *trans*-JA, the most abundant form of the hormone, but levels of *cis*-JA were ∼6 fold higher in aphid-treated plants than control plants (Fig. 2). We also observed that aphid pre-treatment did not have a significant effect on the magnitude or the timing of wounding-induced accumulation of *trans*-JA or *cis*-JA (Fig. S1). We further quantified levels of JA-isoleucine (JA-Ile), the bioactive amino acid conjugate form of the hormone, and found no difference between control and aphid-infested leaves (Fig. 2). These results could indicate that plants respond to aphid feeding by accumulating *cis*-JA, but aphids block activation of the hormone by the jasmonate-amido synthetase JAR1 or prevent accumulation of JA-Ile by activating its catabolism.

**Figure 2.**
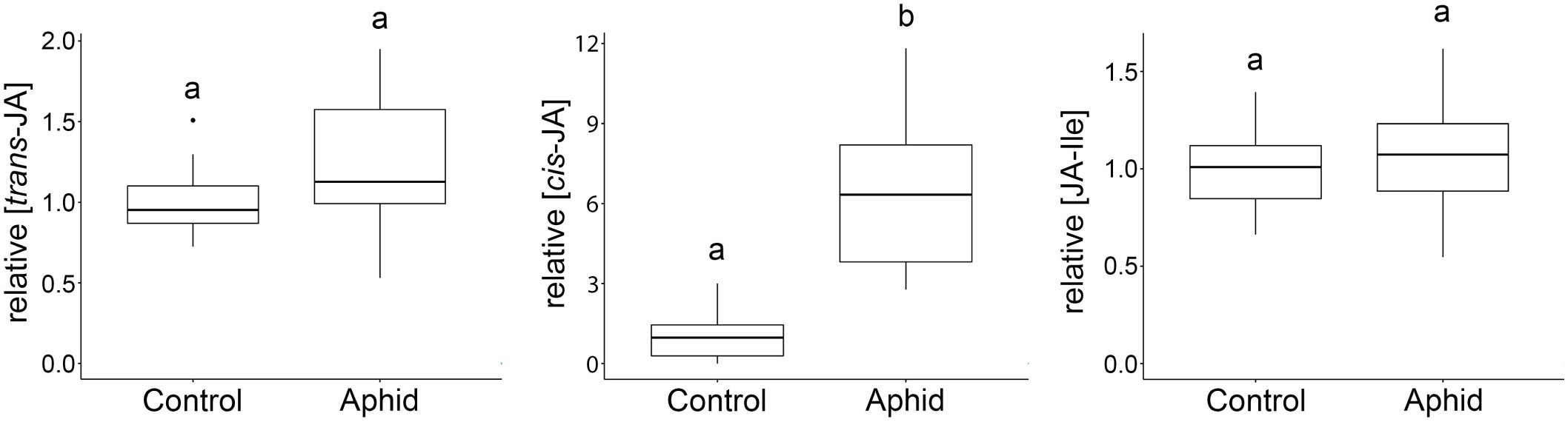
Accumulation of jasmonates in response to aphid feeding. Leaf samples were collected from control plants or plants infested with aphid for seven days. After aphid removal, leaves were extracted and trans-JA (**a**), cis-JA (**b**), and JA-Ile (**c**) were quantified by liquid chromatography– mass spectrometry (LC-MS). Different letters indicate significance at P<0.05 according to two-tailed unpaired t-test. Data from three or four independent experiments were included in the analysis.

To test the latter possibility, we analyzed expression of soybean orthologs of the CYP94C1 protein that catalyze conversion of 12OH-JA-Ile to 12COOH-JA-Ile during JA-Ile degradation (Heitz *et al*. 2012). The two orthologs of CYP94C1 in the soybean genome [*CYP94C1a* (*Glyma.12g087200*) and *CYP94C1b* (*Glyma.11g185700*)] were significantly induced after 7 days of aphid feeding (Fig. 3). However, because several genes involved in JA metabolism are induced with aphid colonization (Studham and MacIntosh 2013), changes in expression of *CYP94C1* orthologs could be part of canonical JA-regulated expression. Thus, we compared expression of the two CYP94C1 orthologs and *PinN2* (as representative of canonical JA-response) in response to mechanical wounding or aphid infestation. Expression of the three genes was induced by both treatments (Fig 3a-c). However, *PinN2* expression was significantly higher in response to wounding than to aphids, while both *CYP94C1* genes were expressed at much higher levels in response to aphids than wounding. Analysis of the ratio aphid/wounding (Fig. 3d) shows that *PinN2* was preferentially expressed in response to wounding, while both JA-Ile catabolism genes were preferentially expressed in response to aphids.

**Figure 3.**
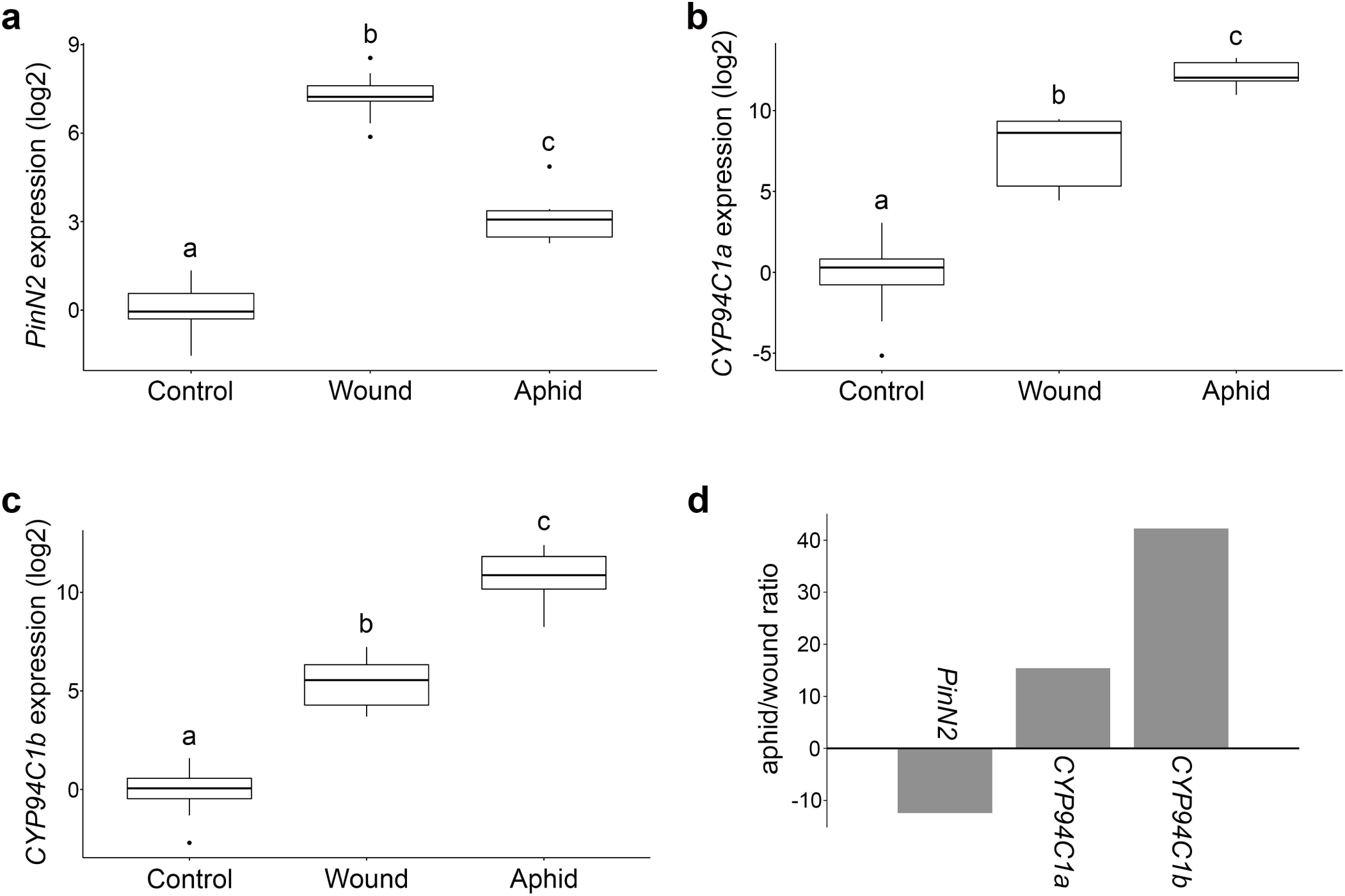
Comparison of the soybean jasmonate response to wounding and aphid infestation. Leaf RNA was extracted from control plants, leaves wounded for six hours, or leaves infested by aphids for seven days. Expression of *PinN2* as marker for JA-dependent defensive output (**a**) or *CYP94C1a* (**b**) and *CYP94C1b* (**c**) as markers for JA-Ile catabolism was quantified by quantitative PCR. Expression was normalized using *UBQ* as internal control. The ratio aphid expression/wounding expression for each gene is depicted in (**d**). Different letters indicate significance at P<0.05 according to two-tailed unpaired t-test. Data from three independent experiments were included in the analysis.

### Exogenous ABA pre-treatment antagonizes wound-induced JA signaling in soybean

We quantified ABA levels and found a significant increase for this hormone (43%) in aphid-infested plants compared to control plants (Fig. 4a and Supporting Table S2). To test whether ABA had a negative effect on JA signaling in soybean, we pre-treated plants with 0µM or 100µM ABA for 24 hours prior to wounding and analyzed JA-dependent responses by quantifying *PinN2* expression (Fig. 4b). In unwounded plants, ABA had a small effect on *PinN2* transcript levels and, as expected, wounding highly induced *PinN2* transcript levels. However, if plants had been previously treated with ABA, they failed to fully induce *PinN2* transcripts in response to wounding, and the pattern was reminiscent of aphid-regulated attenuation of the JA pathway. These results indicate that ABA itself can block JA-regulated responses in soybean and are consistent with a role for ABA as an antagonist of effective JA defenses against aphids.

**Figure 4.**
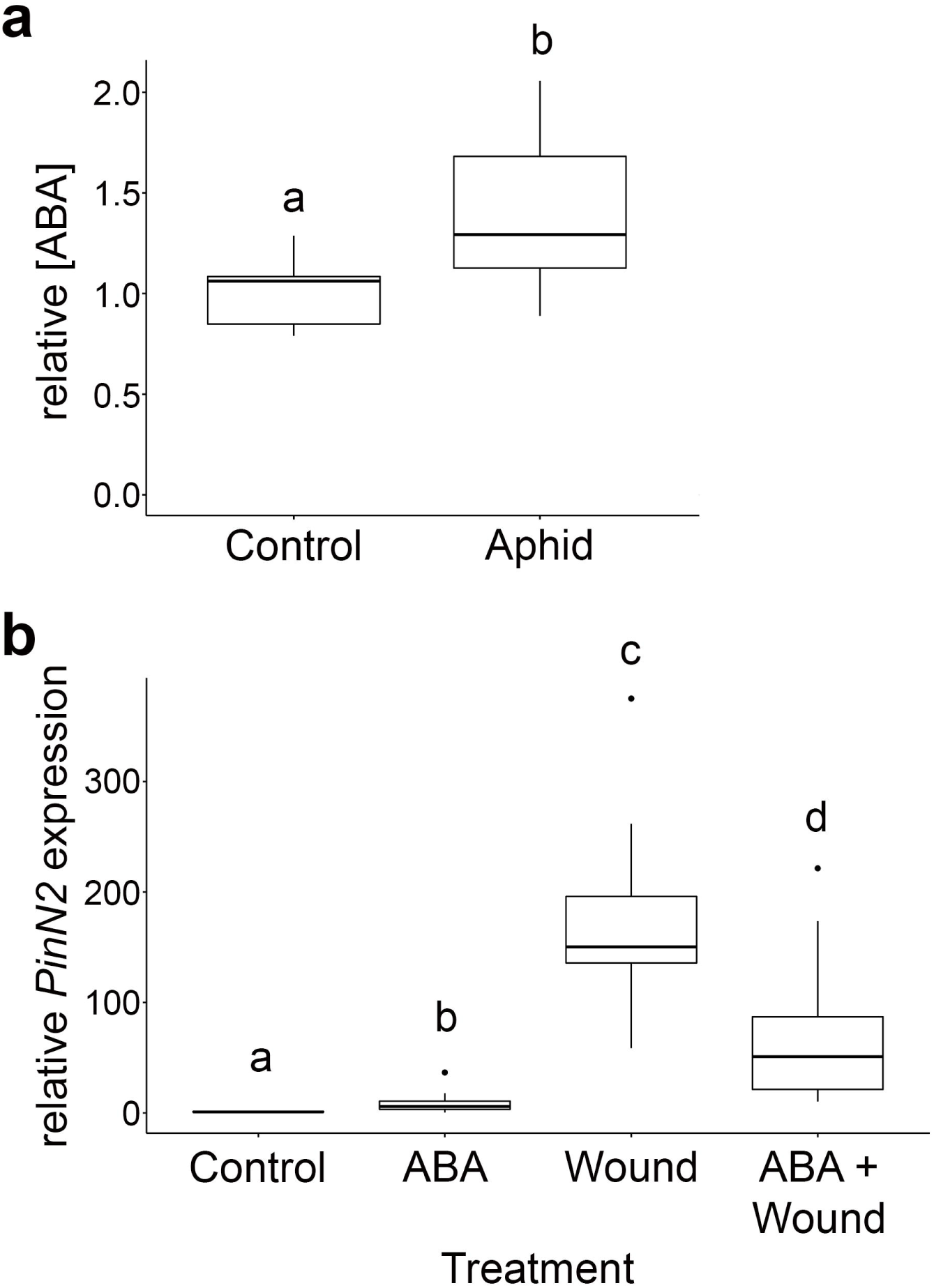
ABA is a candidate antagonist of JA-responses. (**a**) Aphid colonization causes an increase in ABA levels. Leaf samples were collected from control plants or plants infested with aphid for seven days. After aphid removal, leaves were extracted and abscisic acid (ABA) was quantified by liquid chromatography– mass spectrometry (LC-MS). (**b**) Exogenous abscisic acid treatment suppresses wound-induced JA signaling. Plants were exposed to 0µM or 100µM (±ABA) for 24 hours then wounded. Six hours after wounding, samples were collected. Relative expression of *PinN2* was assayed using quantitative PCR. Different letters indicate significance at P<0.05, LSD multiple comparisons test.

### Soybean aphids have compromised performance on soybean plants with defects in the ABA pathway

To determine if ABA is important for population growth of aphids in the compatible interaction, we knocked down expression of genes involved in ABA biosynthesis and signaling using virus-induced gene silencing (VIGS) via a *bean pod mottle virus* (BPMV) system optimized for soybean (Zhang *et al*. 2010; Whitham *et al*. 2016). Our first VIGS target was *ABA2* (*Glyma.11g151400*), which encodes *XANTHOXIN DEHYDROGENASE*, a short-chain dehydrogenase involved in conversion of xanthoxin to abscisic aldehyde during the penultimate step in ABA biosynthesis (González-Guzmán *et al*. 2002). To determine if ABA downstream signaling is important for aphid performance, we also knocked down *SCOF-1* (*Glyma.17g236200*), a cold-and ABA-regulated transcriptional activator that positively regulates ABA-responsive element (ABRE)-binding transcription factors (Kim *et al*. 2001). In susceptible plants, *SCOF-1* expression is highly upregulated by aphid feeding at 1 and 7 days after infestation (4.2 and 10.2 fold, respectively; (Studham and MacIntosh 2013), suggesting that this transcription factor may be a key regulator of the ABA response during soybean aphid infestation of susceptible soybeans. We verified gene knockdown by qRT-PCR; relative to the vector control plants, the transcript level of *ABA2* was reduced 91% in *aba2* RNAi plants while *SCOF-1* was reduced 67% in *scof-1* RNAi plants (Fig. S2). Knockdown of *ABA2* and *SCOF-1* resulted in 24.7% and 25.2% lower aphid populations, respectively, compared to vector controls, and 31.1% and 31.5%, respectively, compared to mock-inoculated plants (Fig. 5). These results indicate that a functional ABA pathway is important for optimal aphid population growth.

**Figure 5.**
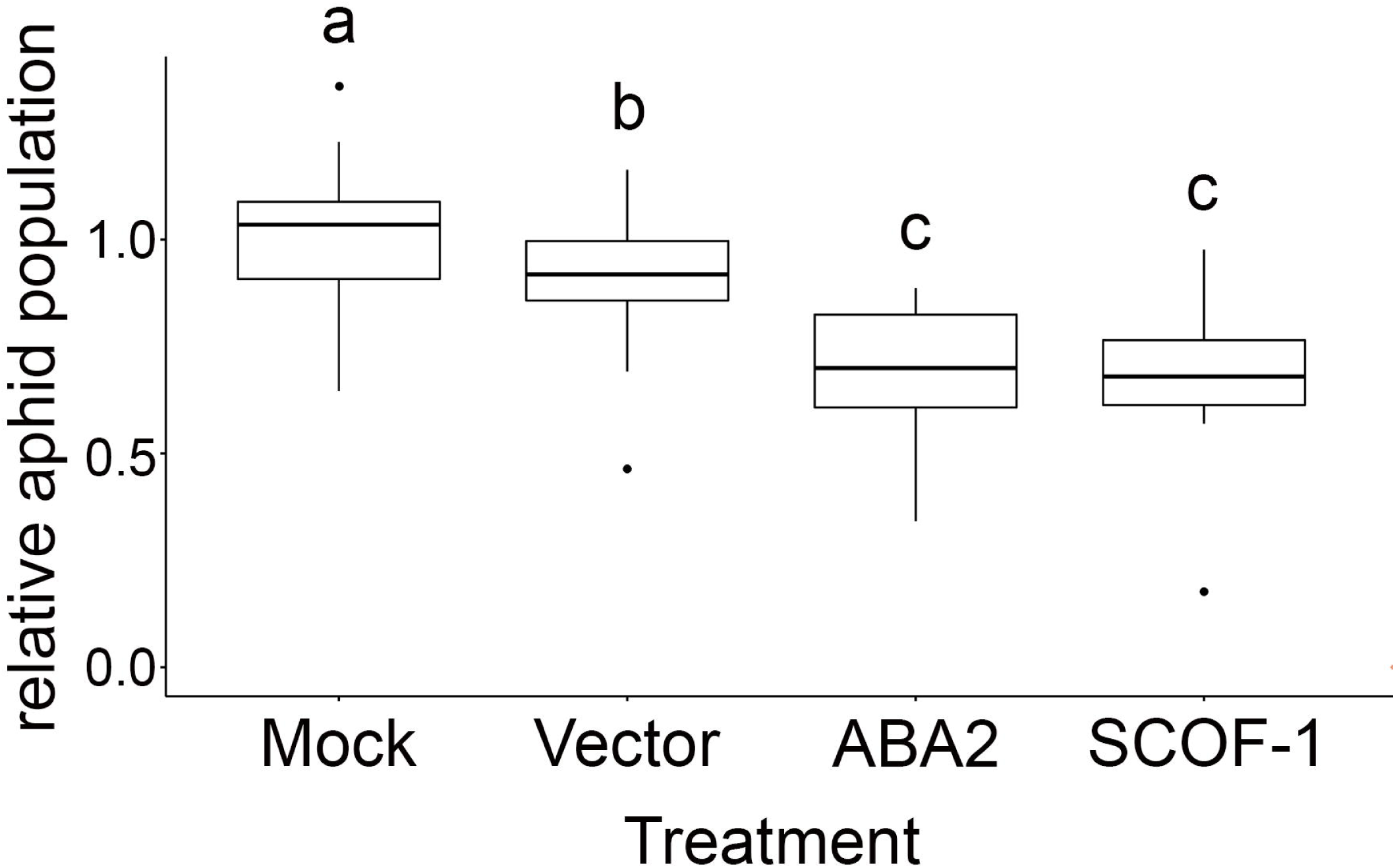
Soybean aphid populations are lower on ABA-knockdown plants. Plants at the V0 growth stage were mock-inoculated, infected with virus-induced gene silencing (VIGS) empty vector, or with vectors targeting *ABA2* or *SCOF-1*. After 20 days, ten age-synchronized apterous aphids were placed on the abaxial surface of a V3 leaf in each plant and confined within clip cages. Aphid population was quantified seven days after infestation. Different letters indicate significance at P<0.05, LSD multiple comparisons test.

### ABA controls basal levels of JA signaling in soybean and aphids exploit this ABA-JA crosstalk

To further investigate ABA-JA crosstalk in soybean, we analyzed basal expression of *PinN2* in control conditions (i.e. free of aphids or wounding stress). We found that *PinN2* was expressed at a higher basal level in both *aba2* RNAi and *scof-1* RNAi plants compared to mock-treated and vector control plants, which were not significantly different from each other (Fig. 6a). Basal *PinN2* levels in *aba2* RNAi and *scof-1* RNAi plants were similar to levels of *PinN2* in wounded mock plants (compare expression levels to those in Fig. 6b, P=0.3734 and P=0.2716 for comparisons against wounded mock plants). Thus, it is clear that defects in ABA biosynthesis or signaling de-repress expression of JA-responsive genes, revealing an antagonistic crosstalk between ABA and JA pathways in soybean. This may be a mechanism used by soybean plants to ensure low levels of JA-responsive gene expression in the absence of stress.

**Figure 6.**
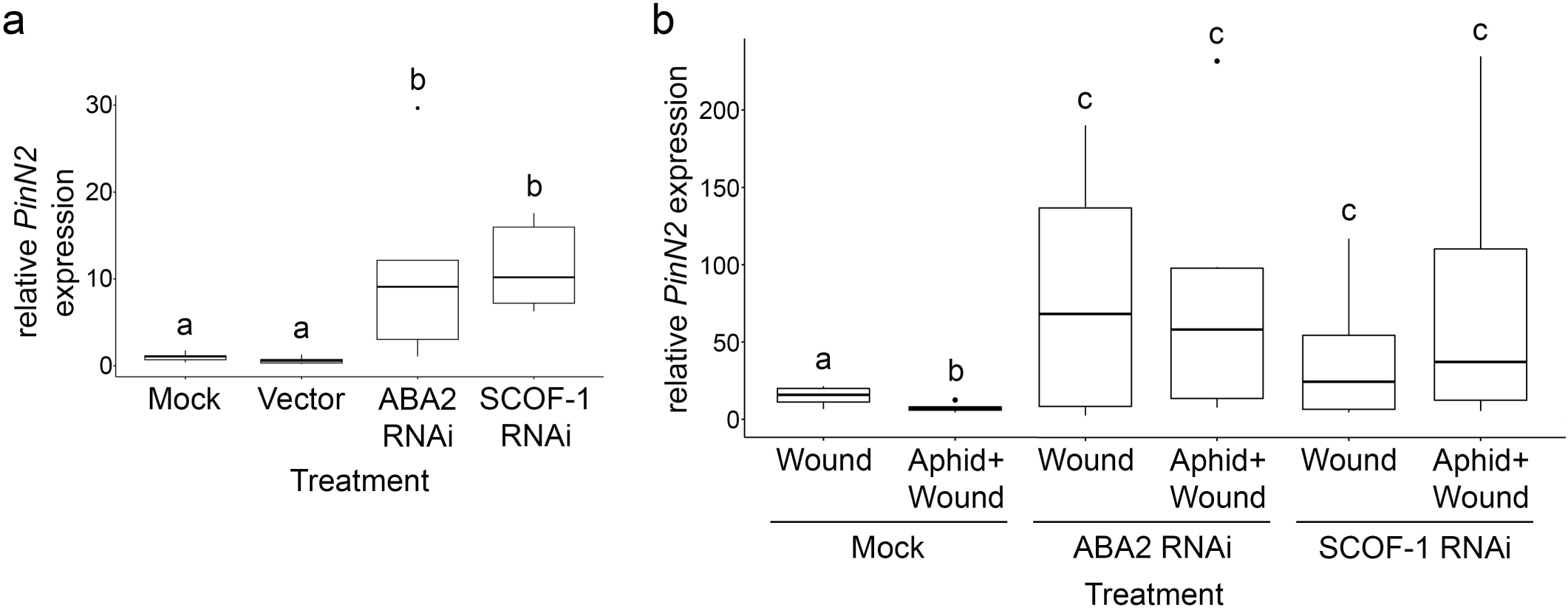
ABA is necessary for suppression of JA signaling. (**a**) Basal JA-dependent gene expression is higher in ABA-knockdown plants. Plants at the V0 growth stage were mock-inoculated, infected with virus-induced gene silencing (VIGS) empty vector, or with vectors targeting *ABA2* or *SCOF-1*. After 20 days, RNA was extracted from V3 leaves of plants grown in the absence of other stressors and expression of *PinN2* was quantified by qPCR. (**b**) Aphid-triggered suppression of JA signaling is mediated through the ABA pathway. Plants were treated with VIGS vectors as in (a). Expression level of *PinN2* was measured using qPCR in wounded plants (W) or wounded plants that had been exposed to aphid feeding for 7 days (AW). Different letters indicate significance at P<0.05, LSD multiple comparisons test.

We then tested whether soybean aphids need the soybean ABA pathway to attenuate wound-induced accumulation of *PinN2* observed in previous experiments. We repeated our first experiment (results presented in in Fig. 1a) using control (mock) plants or plants with knock-down expression of *ABA2* or *SCOF-1*. As found previously (Fig. 1), we confirmed that, compared to plants without aphids, previous exposure of non-silenced plants to aphid feeding attenuated wound induction of *PinN2* (Fig. 6b). However, in *aba2* or *scof-1* RNAi plants, aphid pre-treatment did not attenuate accumulation of *PinN2* in response to wounding. Moreover, we observed significant increases in the content of JA-Ile in aphid-infested *aba2* RNAi plants compared to aphid-infested mock or vector plants (Figure S3). Thus, defects in the ABA pathway prevented aphids from interfering with JA defenses. These data support the hypothesis that soybean aphids exploit an ABA-JA antagonism to modulate plants defenses and increase the colonization of their host.

## Discussion

Feeding by soybean aphids suppresses host-plant defenses and improves the performance of conspecifics. Plants previously colonized by aphids (inducer population) can increase the performance of subsequent aphid populations (response population; (Varenhorst *et al*. 2015b). This effect is dependent on the density of inducer populations (Varenhorst *et al*. 2015b) and the effects remain for at least 24 h after removal of inducer populations (Varenhorst *et al*. 2015a). Both virulent and avirulent inducer populations can increase performance of response populations through two proposed mechanisms, feeding facilitation and obviation of resistance (Neupane *et al*. 2019; Varenhorst *et al*. 2015a; Varenhorst *et al*. 2015b). In addition, aphid colonization can facilitate colonization by soybean cyst nematodes (McCarville *et al*. 2012) and facilitate feeding by lepidopteran larvae (Fig. 1).

Suppression of JA-dependent defenses can explain, at least in part, these effects. Previous studies have shown that JA signals induce effective defenses against soybean aphids. Treatment of susceptible soybean plants with MeJA 24 h before infestation with virulent aphids resulted in a 30% reduction in growth of soybean aphid populations (Selig *et al*. 2016). Similarly, JA-Ile treatment (6 h before infestation) resulted in drastic reductions of populations of virulent aphids on soybean plants (Yates-Stewart *et al*. 2020). Interestingly, SA does not seem to interfere with JA-dependent defenses against soybean aphids. Simultaneous treatment with MeJA and SA has a similar effect on soybean aphid performance as the MeJA-only treatment (Selig *et al*. 2016), and SA treatment increased resistance of a soybean line carrying the *Rag1* resistance gene (Studham and MacIntosh 2013). Additionally, our previous transcriptome analysis (Studham and MacIntosh 2013) indicated that, at early stages of infestation, both SA and JA biosynthesis and signaling are activated in parallel, while the two hormone signals seem to be repressed after 7 days of infestation. These results suggested that susceptible plants defend against soybean aphids using JA-dependent defenses, but these defenses are not effectively deployed in compatible interactions, likely due to an induced-susceptibility mechanism (Varenhorst *et al*. 2015b).

Here, we showed that aphids reduce plant defenses by suppressing JA signals. Previous analysis of changes in content of polyunsaturated fatty acids initially led us to suggest that this block in JA-dependent defenses could be due to a reduction in the ability of plants to produce JA (Kanobe *et al*. 2015). However, we found that soybean plants previously colonized by aphids were not able to respond to JA treatment as effectively as control plants, indicating that the block in JA-signaling takes place downstream of JA synthesis. Supporting this finding, aphid-infested plants accumulate *cis*-JA, but levels of JA-Ile do not change with aphid feeding. Moreover, we observed a preferential increase in expression of genes encoding CYP94C1, an enzyme that specifically participates in JA-Ile catabolism. During catabolism, JA-Ile is oxidized by JA-Ile o-hydroxylases CYP94B3 and B1 that generate 12OH-JA-Ile (Heitz *et al*. 2012; Widemann *et al*. 2015). Subsequently, CYP94C1 catalyzes conversion of 12OH-JA-Ile to 12COOH-JA (Heitz *et al*. 2012; Widemann *et al*. 2015). Arabidopsis plants overexpressing *CYP94C1* show a decrease in accumulation of JA-Ile after wounding (Heitz *et al*. 2012). Similarly, CYP94C1-OE plants are more susceptible to the fungal pathogen *Botrytis cinerea* due to their inability to activate JA-dependent responses (Aubert *et al*. 2015). Our results suggest that aphid interference with JA signaling is due, at least in part, to a differential induction of JA catabolism by aphid feeding.

Recently, a similar mechanism for modulation of JA catabolism via herbivore feeding was proposed for interactions between spider mites and Arabidopsis. Spider mite infestation induces expression of *JAO2*, which encodes the enzyme responsible for hydroxylation of JA into 12-OH-JA (Smirnova *et al*. 2017; Rosa-Diaz *et al*. 2023). Spider mite feeding and reproduction were lower in the Arabidopsis *jao2* mutant (Rosa-Diaz *et al*. 2023) indicating that JAO2 acts as a negative regulator of defenses against spider mites by increasing JA catabolism. Several pathogen and herbivore effectors that directly interfere with JA-Ile accumulation or signal transduction have also been identified (Zhang *et al*. 2017; Chen *et al*. 2019; Wu and Ye 2020). This indicates that different pest species have evolved a variety of mechanisms to suppress plant defense by altering defense signaling, and soybean aphids may also use salivary effectors to activate ABA-JA antagonism.

Based on our previous results, we hypothesized that the ability of soybean aphid to block JA responses requires ABA signaling. Here we showed that aphid feeding induces accumulation of ABA, and that ABA is necessary for aphid-triggered attenuation of JA-mediated soybean defenses. Moreover, plants rendered defective in ABA signaling by silencing an ABA biosynthetic gene (*ABA2*) or a key ABA transcriptional regulator (*SCOF-1*) were more resistant to aphid colonization than control plants. Importantly, on these ABA-defective plants, aphids cannot suppress JA-regulated induction of *PinN2*. Thus, in soybean, ABA antagonizes JA-dependent defense outputs, and soybean aphids exploit this antagonism to promote their feeding and colonization.

It was recently shown that ABA-JA antagonism also regulates isoflavone production in soybean (Lin *et al*. 2025). Importantly, we have previously shown that soybean isoflavones have a deterrent effect on soybean aphid feeding (Hohenstein *et al*. 2019). ABA negatively regulates isoflavone biosynthesis (Ward *et al*. 1989; Mohr and Cahill 2001; Graham and Graham 1996), while JA could be a positive regulator of the pathway (Graham and Graham 1996), although other oxylipins are more effective elicitors (Fliegmann *et al*. 2003). Transcriptome data mined to identify regulators of isoflavone production indicated that JA signaling positively regulates synthesis of these phytoalexins (Lin *et al*. 2025). ABA induces expression of *GmJAZ1* genes (1-9) that encode jasmonate-zim domain (JAZ) transcriptional repressors (Lin *et al*. 2025) and these JAZ1 repressors negatively regulate expression of glyceollin biosynthetic genes in soybeans through a direct interaction with the transcription factor GmNAC42-1. GmNAC42-1 is one of the key transcription factor activators of the isoflavone pathway (Jahan *et al*. 2019); binding of JAZ1 repressor to the transcription factor inhibits the ability of GmNAC42-1 to transactivate glyceollin biosynthesis gene promoters (Lin *et al*. 2025). Together these findings suggest that ABA antagonism of JA signaling may also play a role in suppressing isoflavone defenses that contribute to increased plant susceptibility to soybean aphids.

ABA has historically been associated with water stress responses in plants (Bray 2002). In soybean, drought stress exacerbates soybean aphid populations in the field (Rice *et al*. 2004). Moreover, higher soybean aphid populations grew on drought-stressed soybean plants than on plants maintained in overwatered soil conditions, a result attributed to higher ABA levels in drought-stressed plants and suppression of JA and SA signaling (Nachappa *et al*. 2016). Dehydration stress also upregulates JAZ repressors in soybean plants (Lin *et al*. 2025), further supporting the role of ABA in antagonizing JA signaling. Intriguingly, compared to susceptible plants, soybean plants that are tolerant to aphid feeding have constitutively elevated ABA and JA levels and gene expression related to these hormones (Chapman *et al*. 2018).

JA-ABA antagonism has been studied in other plant species, and their interaction seems to be complex and context-dependent. ABA treatments antagonized JA-dependent defense against migratory nematodes (Nahar *et al*. 2012) and black-streaked dwarf virus (Xie *et al*. 2018) in rice. In Arabidopsis, ABA activated the MYC2-branch of JA signaling (with *VSP1* as a marker gene) but repressed the ethylene-modulated (ERF/ORA59) branch of JA signaling (with *PDF1.2* as a marker gene (Anderson *et al*. 2004; Kazan and Manners 2013; Vos *et al*. 2019). Thus, in Arabidopsis, ABA activation of the MYC-branch was necessary for effective defense against herbivores (Vos *et al*. 2019). In soybean, however, we observed increases in *PinN2* expression in ABA defective plants, indicating that in this plant species ABA was antagonistic to herbivore defenses. We can speculate that ABA induction by soybean aphids may repress the ERF branch of JA signaling, and that rewiring of JA-signaling pathways in soybean resulted in the recruitment of the JA-ET branch to promote herbivore defenses. Consistent with this idea, soybean *PinN2* is responsive to wounding only in the presence of a functional ethylene signaling pathway (Botella *et al*. 1996).

We found that soybean plants with defects in ABA biosynthesis or signaling had much higher levels of JA signaling in the absence of stress. In fact, *PinN2* expression in ABA2-RNAi and SCOF-1-RNAi plants without aphid-or wounding-related stressors was equivalent to expression of the gene after wounding in mock or vector only controls. These results indicated that ABA is an important regulator of basal JA signaling in soybean in the absence of stress, and that aphids have coopted this mechanism to promote their feeding and colonization. Induction of ABA signaling to antagonize JA is probably one of the mechanisms used by aphids to suppress defenses in susceptible soybean; however, it is likely just one facet of their overall feeding and colonization strategy. ABA-deficient plants have lower aphid populations than susceptible controls, but those are still larger than what has been observed in resistant *Rag* plants. Our transcriptome data also suggest that SA is repressed in susceptible plants (Studham and MacIntosh 2013). Whether ABA is also able to antagonize SA signaling in soybean needs to be tested. Follow up transcriptome studies and targeted approaches to silence other hormone pathways will help us dissect crosstalk mechanisms among hormones involved in soybean defense and susceptibility to soybean aphid.

## Acknowledgements

We thank Dr. Steve Whitham (Iowa State University) and Dr. Michelle Graham (USDA-ARS) for providing vectors and helping set up VIGS experiments. We also thank Dr. Russell Jurenka for providing corn earworm larvae for our experiments and for sharing knowledge on caterpillar husbandry. This work was funded by a grant from the U.S. Department of Agriculture (USDA) AFRI/NIFA program (# 2019-67013-29352) to G.C.M.

## Competing interests

None declared.

## Author contributions

J.D.H., C.K., and G.C.M. conceived the project and designed experiments. J.D.H., C.K., M.I.N., P.G., D.Z., N.K., A.M.H., and J.F.T. performed experiments and analyzed data. J.D.H., C.K., and G.C.M. prepared the manuscript. All authors revised the manuscript.

## Data availability

All generated and analyzed data from this study are included in the published article and its Supporting Information. Correspondence and requests for materials should be addressed to G.C.M.

## Supporting Information

**Figure S1.** Accumulation of jasmonates in response to wounding after aphid feeding.

**Figure S2.** VIGS knockdown confirmation.

**Figure S3.** Accumulation of JA-Ile in response to aphid feeding in control and ABA-deficient plants.

**Table S1**. Primers used to generate RNAi constructs and quantify gene expression (qRT-PCR).

**Table S2**. Average (± SEM) phytohormone content in control and aphid-infested plants 7 days after infestation.

## Notes

### Competing Interest Statement

The authors have declared no competing interest.

## References

Åhman I, Kim S-Y, Zhu L-H. 2019. Plant Genes Benefitting Aphids—Potential for Exploitation in Resistance Breeding. Frontiers in Plant Science 10: 1452.

Ali JG, Agrawal AA. 2014. Asymmetry of plant-mediated interactions between specialist aphids and caterpillars on two milkweeds. Functional Ecology 28: 1404–1412.

Anderson JP, Badruzsaufari E, Schenk PM, Manners JM, Desmond OJ, Ehlert C, Maclean DJ, Ebert PR, Kazan K. 2004. Antagonistic interaction between abscisic acid and jasmonate-ethylene signaling pathways modulates defense gene expression and disease resistance in Arabidopsis. Plant Cell 16:3460–3479.

Aubert Y, Widemann E, Miesch L, Pinot F, Heitz T. 2015. CYP94-mediated jasmonoyl-isoleucine hormone oxidation shapes jasmonate profiles and attenuates defence responses to *Botrytis cinerea* infection. Journal of Experimental Botany 66:3879–3892.

Botella MA, Xu Y, Prabha TN, Zhao Y, Narasimhan ML, Wilson KA, Nielsen SS, Bressan RA, Hasegawa PM. 1996. Differential Expression of Soybean Cysteine Proteinase Inhibitor Genes during Development and in Response to Wounding and Methyl Jasmonate. Plant Physiology 112:1201–1210.

Bray EA. 2002. Abscisic acid regulation of gene expression during water-deficit stress in the era of the *Arabidopsis* genome. *Plant*, Cell & Environment 25:153–161.

Cao H-H, Wang S-H, Liu T-X. 2014. Jasmonate-and salicylate-induced defenses in wheat affect host preference and probing behavior but not performance of the grain aphid, *Sitobion avenae*. Insect Science 21:47–55.

Chapman KM, Marchi-Werle L, Hunt TE, Heng-Moss TM, Louis J. 2018. Abscisic and Jasmonic Acids Contribute to Soybean Tolerance to the Soybean Aphid (*Aphis glycines* Matsumura). Scientific Reports 8:15148.

Chen C-Y, Liu Y-Q, Song W-M, Chen D-Y, Chen F-Y, Chen X-Y, Chen Z-W, Ge S-X, Wang C-Z, Zhan S, Chen X-Y, Mao Y-B. 2019. An effector from cotton bollworm oral secretion impairs host plant defense signaling. Proceedings of the National Academy of Sciences 116:14331–14338.

Cooper WR, Goggin FL. 2005. Effects of jasmonate-induced defenses in tomato on the potato aphid, *Macrosiphum euphorbiae*. Entomologia Experimentalis et Applicata 115:107–115.

Ellis C, Karafyllidis L, Turner JG. 2002. Constitutive activation of jasmonate signaling in an *Arabidopsis* mutant correlates with enhanced resistance to *Erysiphe cichoracearum*, *Pseudomonas syringae*, and *Myzus persicae*. Molecular Plant-Microbe Interactions 15:1025–1030.

Erb M, Reymond P. 2019. Molecular Interactions Between Plants and Insect Herbivores. Annual Review of Plant Biology 70:527–557.

Fehr WR, Caviness CE. 1977. STAGES OF SOYBEAN DEVELOPMENT. Iowa Agricultural and Home Economics Experiment Station Special Report 80:3–11

Fliegmann J, Schüler G, Boland W, Ebel J, Mithöfer A. 2003. The Role of Octadecanoids and Functional Mimics in Soybean Defense Responses. Biological Chemistry 384:437–446.

Forcat S, Bennett MH, Mansfield JW, Grant MR. 2008. A rapid and robust method for simultaneously measuring changes in the phytohormones ABA, JA and SA in plants following biotic and abiotic stress. Plant Methods 4:16.

González-Guzmán M, Apostolova N, Bellés JM, Barrero JM, Piqueras P, Ponce MR, Micol JL, Serrano R, Rodríguez PL. 2002. The Short-Chain Alcohol Dehydrogenase ABA2 Catalyzes the Conversion of Xanthoxin to Abscisic Aldehyde. The Plant Cell 14:1833–1846.

Graham TL, Graham MY. 1996. Signaling in Soybean Phenylpropanoid Responses (Dissection of Primary, Secondary, and Conditioning Effects of Light, Wounding, and Elicitor Treatments). Plant Physiology 110:1123–1133.

Heitz T, Widemann E, Lugan R, Miesch L, Ullmann P, Désaubry L, Holder E, Grausem B, Kandel S, Miesch M, Werck-Reichhart D, Pinot F. 2012. Cytochromes P450 CYP94C1 and CYP94B3 Catalyze Two Successive Oxidation Steps of Plant Hormone Jasmonoyl-isoleucine for Catabolic Turnover. Journal of Biological Chemistry 287:6296–6306.

Hillwig MS, Chiozza M, Casteel CL, Lau ST, Hohenstein J, Hernández E, Jander G, MacIntosh GC. 2016. Abscisic acid deficiency increases defence responses against *Myzus persicae* in Arabidopsis. Molecular Plant Pathology 17:225–235.

Hogenhout SA, Bos JIB. 2011. Effector proteins that modulate plant–insect interactions. Current Opinion in Plant Biology 14:422–428.

Hohenstein JD, Studham ME, Klein A, Kovinich N, Barry K, Lee Y-J, MacIntosh GC. 2019. Transcriptional and Chemical Changes in Soybean Leaves in Response to Long-Term Aphid Colonization. Frontiers in Plant Science 10:310.

Jahan MA, Harris B, Lowery M, Coburn K, Infante AM, Percifield RJ, Ammer AG, Kovinich N. 2019. The NAC family transcription factor GmNAC42–1 regulates biosynthesis of the anticancer and neuroprotective glyceollins in soybean. BMC Genomics 20:149

Kaloshian I, Walling LL. 2016. Plant Immunity: Connecting the Dots Between Microbial and Hemipteran Immune Responses. In: Czosnek H, Ghanim M, eds. Management of Insect Pests to Agriculture: Lessons Learned from Deciphering their Genome, Transcriptome and Proteome. Cham, Switzerland: Springer International Publishing, 217–243.

Kanobe C, McCarville MT, O’Neal ME, Tylka GL, MacIntosh GC. 2015. Soybean Aphid Infestation Induces Changes in Fatty Acid Metabolism in Soybean. PLoS One 10:e0145660.

Kazan K, Manners JM. 2013. MYC2: The Master in Action. Molecular Plant 6:686–703.

Kerchev PI, Karpińska B, Morris JA, Hussain A, Verrall SR, Hedley PE, Fenton B, Foyer CH, Hancock RD. 2013. Vitamin C and the abscisic acid-insensitive 4 transcription factor are important determinants of aphid resistance in *Arabidopsis*. Antioxidants & Redox Signaling 18:2091–2105.

Kersch-Becker MF, Thaler JS. 2019. Constitutive and herbivore-induced plant defences regulate herbivore population processes. Journal of Animal Ecology 88:1079–1088.

Kim JC, Lee SH, Cheong YH, Yoo C-M, Lee SI, Chun HJ, Yun D-J, Hong JC, Lee SY, Lim CO, Cho MJ. 2001. A novel cold-inducible zinc finger protein from soybean, SCOF-1, enhances cold tolerance in transgenic plants. The Plant Journal 25:247–259.

Koramutla MK, Kaur A, Negi M, Venkatachalam P, Bhattacharya R. 2014. Elicitation of jasmonate-mediated host defense in *Brassica juncea* (L.) attenuates population growth of mustard aphid *Lipaphis erysimi* (Kalt.). Planta 240:177–194.

Kovinich N, Saleem A, Arnason JT, Miki B. 2011. Combined analysis of transcriptome and metabolite data reveals extensive differences between black and brown nearly-isogenic soybean (*Glycine max*) seed coats enabling the identification of pigment isogenes. BMC Genomics 12:381.

Libault M, Wan J, Czechowski T, Udvardi M, Stacey G. 2007. Identification of 118 Arabidopsis transcription factor and 30 ubiquitin-ligase genes responding to chitin, a plant-defense elicitor. Molecular Plant-Microbe Interactions 20:900–911.

Lin J, Monsalvo I, Jahan MA, Ly M, Wi D, Martirosyan I, Jahan I, Kovinich N. 2025. ABA-regulated JAZ1 suppresses phytoalexin biosynthesis by binding GmNAC42-1 in soybean. Current Plant Biology 42:100453.

Losvik A, Beste L, Glinwood R, Ivarson E, Stephens J, Zhu L-H, Jonsson L. 2017. Overexpression and Down-Regulation of Barley Lipoxygenase LOX2.2 Affects Jasmonate-Regulated Genes and Aphid Fecundity. International Journal of Molecular Sciences 18:2765.

Ma K, Tang Q, Liang P, Xia J, Zhang B, Gao X. 2019. Toxicity and sublethal effects of two plant allelochemicals on the demographical traits of cotton aphid, *Aphis gossypii* Glover (Hemiptera: Aphididae). PLoS One 14:e0221646.

McCarville MT, O’Neal M, Tylka GL, Kanobe C, MacIntosh GC. 2012. A nematode, fungus, and aphid interact via a shared host plant: implications for soybean management. Entomologia Experimentalis et Applicata 143:55–66.

Mewis I, Appel HM, Hom A, Raina R, Schultz JC. 2005. Major signaling pathways modulate Arabidopsis glucosinolate accumulation and response to both phloem-feeding and chewing insects. Plant Physiology 138:1149–1162.

Mewis I, Tokuhisa JG, Schultz JC, Appel HM, Ulrichs C, Gershenzon J. 2006. Gene expression and glucosinolate accumulation in *Arabidopsis thaliana* in response to generalist and specialist herbivores of different feeding guilds and the role of defense signaling pathways. Phytochemistry 67:2450–2462.

Mohr PG, Cahill DM. 2001. Relative roles of glyceollin, lignin and the hypersensitive response and the influence of ABA in compatible and incompatible interactions of soybeans with *Phytophthora sojae*. Physiological and Molecular Plant Pathology 58:31–41.

Nachappa P, Culkin CT, Saya PM, Han J, Nalam VJ. 2016. Water Stress Modulates Soybean Aphid Performance, Feeding Behavior, and Virus Transmission in Soybean. Frontiers in Plant Science 7:552.

Nahar K, Kyndt T, Nzogela YB, Gheysen G. 2012. Abscisic acid interacts antagonistically with classical defense pathways in rice–migratory nematode interaction. New Phytologist 196:901–913.

Natukunda MI, MacIntosh GC. 2020. The Resistant Soybean-*Aphis glycines* Interaction: Current Knowledge and Prospects. Frontiers in Plant Science 11:1223.

Neupane S, Purintun JM, Mathew FM, Varenhorst AJ, Nepal MP. 2019. Molecular Basis of Soybean Resistance to Soybean Aphids and Soybean Cyst Nematodes. Plants 8:374.

Paz MM, Martinez JC, Kalvig AB, Fonger TM, Wang K. 2006. Improved cotyledonary node method using an alternative explant derived from mature seed for efficient *Agrobacterium*-mediated soybean transformation. Plant Cell Reports 25:206–213.

Rice ME, O’Neal ME, Pedersen P. 2004. Soybean Aphids in Iowa-2004. Iowa State University SP 247. https://dr.lib.iastate.edu/handle/20.500.12876/33225

Rosa-Diaz I, Santamaria ME, Acien JM, Diaz I. 2023. Jasmonic acid catabolism in Arabidopsis defence against mites. Plant Science 334:111784.

Schaeffer RN, Wang Z, Thornber CS, Preisser EL, Orians CM. 2018. Two invasive herbivores on a shared host: patterns and consequences of phytohormone induction. Oecologia 186:973–982.

Schmelz EA, Engelberth J, Tumlinson JH, Block A, Alborn HT. 2004. The use of vapor phase extraction in metabolic profiling of phytohormones and other metabolites. The Plant Journal 39:790–808.

Selig P, Keough S, Nalam VJ, Nachappa P. 2016. Jasmonate-dependent plant defenses mediate soybean thrips and soybean aphid performance on soybean. Arthropod-Plant Interactions 10:273–282.

Smirnova E, Marquis V, Poirier L, Aubert Y, Zumsteg J, Ménard R, Miesch L, Heitz T. 2017. Jasmonic Acid Oxidase 2 Hydroxylates Jasmonic Acid and Represses Basal Defense and Resistance Responses against *Botrytis cinerea* Infection. Molecular Plant 10:1159–1173.

Soler R, Badenes-Pérez FR, Broekgaarden C, Zheng S-J, David A, Boland W, Dicke M. 2012. Plant-mediated facilitation between a leaf-feeding and a phloem-feeding insect in a brassicaceous plant: from insect performance to gene transcription. Functional Ecology 26:156–166.

Studham ME, MacIntosh GC. 2013. Multiple phytohormone signals control the transcriptional response to soybean aphid infestation in susceptible and resistant soybean plants. Molecular Plant-Microbe Interactions 26:116–129.

Takemoto H, Uefune M, Ozawa R, Arimura G-I, Takabayashi J. 2013. Previous infestation of pea aphids *Acyrthosiphon pisum* on broad bean plants resulted in the increased performance of conspecific nymphs on the plants. Journal of Plant Interactions 8:370–374.

Thaler JS, Fidantsef AL, Duffey SS, Bostock RM. 1999. Trade-Offs in Plant Defense Against Pathogens and Herbivores: A Field Demonstration of Chemical Elicitors of Induced Resistance. Journal of Chemical Ecology 25:1597–1609.

Tilmon KJ, Hodgson EW, O’Neal ME, Ragsdale DW. 2011. Biology of the Soybean Aphid, *Aphis glycines* (Hemiptera: Aphididae) in the United States. Journal of Integrated Pest Management 2:A1–A7.

Varenhorst A, McCarville M, O’Neal M. 2015a. Determining the duration of *Aphis glycines* (Hemiptera: Aphididae) induced susceptibility effect in soybean. Arthropod-Plant Interactions 9:457–464.

Varenhorst AJ, McCarville MT, O’Neal ME. 2015b. An Induced Susceptibility Response in Soybean Promotes Avirulent *Aphis glycines* (Hemiptera: Aphididae) Populations on Resistant Soybean. Environmental Entomology 44:658–667.

Vos IA, Verhage A, Watt LG, Vlaardingerbroek I, Schuurink RC, Pieterse CM, Van Wees SC. 2019. Abscisic acid is essential for rewiring of jasmonic acid-dependent defenses during herbivory. bioRxiv:747345. doi: 10.1101/747345

Walling LL. 2008. Avoiding effective defenses: Strategies employed by phloem-feeding insects. Plant Physiology 146:859–866.

Ward EWB, Cahill DM, Bhattacharyya MK. 1989. Abscisic Acid Suppression of Phenylalanine Ammonia-Lyase Activity and mRNA, and Resistance of Soybeans to *Phytophthora megasperma* f.sp. *glycinea*. Plant Physiology 91:23–27.

Whitham SA, Lincoln LM, Chowda-Reddy RV, Dittman JD, O’Rourke JA, Graham MA. 2016. Virus-Induced Gene Silencing and Transient Gene Expression in Soybean (*Glycine max*) Using Bean Pod Mottle Virus Infectious Clones. Current Protocols in Plant Biology 1: 263–283.

Widemann E, Grausem B, Renault H, Pineau E, Heinrich C, Lugan R, Ullmann P, Miesch L, Aubert Y, Miesch M, Heitz T, Pinot F. 2015. Sequential oxidation of Jasmonoyl-Phenylalanine and Jasmonoyl-Isoleucine by multiple cytochrome P450 of the CYP94 family through newly identified aldehyde intermediates. Phytochemistry 117:388–399.

Will T, van Bel AJE. 2006. Physical and chemical interactions between aphids and plants. Journal of Experimental Botany 57:729–737.

Wu X, Ye J. 2020. Manipulation of Jasmonate Signaling by Plant Viruses and Their Insect Vectors. Viruses 12:148.

Xie K, Li L, Zhang H, Wang R, Tan X, He Y, Hong G, Li J, Ming F, Yao X, Yan F, Sun Z, Chen J. 2018. Abscisic acid negatively modulates plant defence against rice black-streaked dwarf virus infection by suppressing the jasmonate pathway and regulating reactive oxygen species levels in rice. Plant, Cell & Environment 41:2504–2514.

Yao L, Yang B, Ma X, Wang S, Guan Z, Wang B, Jiang Y. 2020. A Genome-Wide View of Transcriptional Responses during *Aphis glycines* Infestation in Soybean. International Journal of Molecular Sciences 21:5191.

Yates-Stewart AD, Pekarcik A, Michel A, Blakeslee JJ. 2020. Jasmonic Acid-Isoleucine (JA-Ile) Is Involved in the Host-Plant Resistance Mechanism Against the Soybean Aphid (Hemiptera: Aphididae). Journal of Economic Entomology 113: 2972–2978.

Zarate SI, Kempema LA, Walling LL. 2007. Silverleaf whitefly induces salicylic acid Defenses and suppresses effectual jasmonic acid defenses. Plant Physiology 143:866–875.

Zhang C, Bradshaw JD, Whitham SA, Hill JH. 2010. The Development of an Efficient Multipurpose Bean Pod Mottle Virus Viral Vector Set for Foreign Gene Expression and RNA Silencing. Plant Physiology 153:52–65.

Zhang L, Zhang F, Melotto M, Yao J, He SY. 2017. Jasmonate signaling and manipulation by pathogens and insects. Journal of Experimental Botany 68:1371–1385.

Zhang P-J, Huang F, Zhang J-M, Wei J-N, Lu Y-B. 2015. The mealybug *Phenacoccus solenopsis* suppresses plant defense responses by manipulating JA-SA crosstalk. Scientific Reports 5:9354.

Zhang P, Zhu X, Huang F, Liu Y, Zhang J, Lu Y, Ruan Y. 2011. Suppression of Jasmonic Acid-Dependent Defense in Cotton Plant by the Mealybug *Phenacoccus solenopsis*. PLoS One 6:e22378.

Zhu-Salzman K, Salzman RA, Ahn J-E, Koiwa H. 2004. Transcriptional regulation of sorghum defense determinants against a phloem-feeding aphid. Plant Physiology 134:420–431.

