## Supplementary Material for "Soybean aphids exploit abscisic acid signaling to suppress jasmonate defense responses"

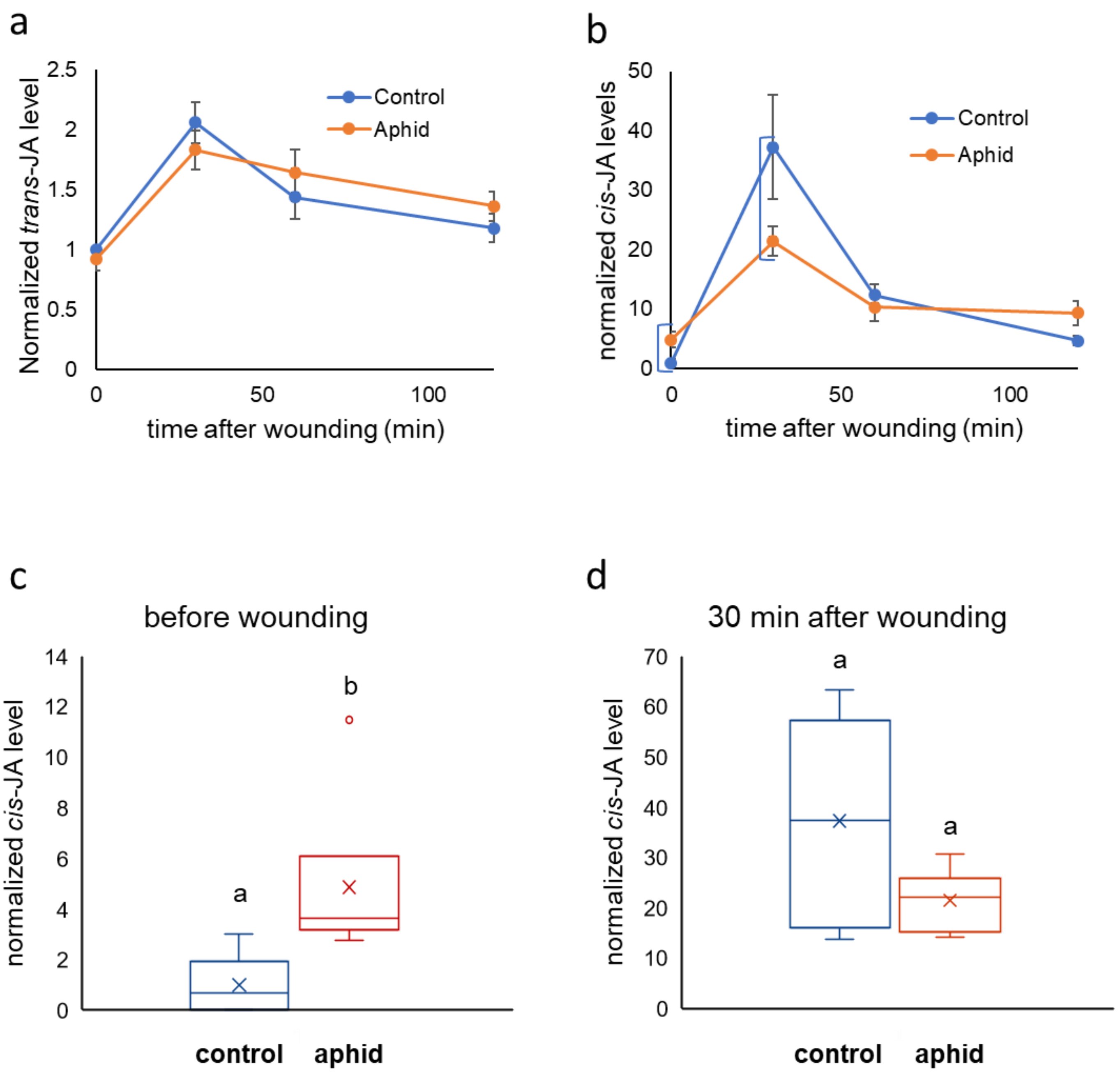


**Fig. S1. Accumulation of jasmonates in response to wounding after aphid feeding.** Plants were exposed to aphids for 7 days (Aphid) or were left unexposed (Control). Plants were then wounded with tweezers and leaf samples were collected at the indicated time points. After aphid removal, leaves were extracted and trans-JA (**a**) and cis-JA (**b**) were quantified by liquid chromatography– mass spectrometry (LC-MS). Levels of *cis*-JA before (**c**) or 30 minutes after wounding (**d**) are shown as box plots. The data are the same as in (b). Different letters indicate significance at P<0.05 according to two-tailed unpaired t-test. Data from two independent experiments were included in the analysis.

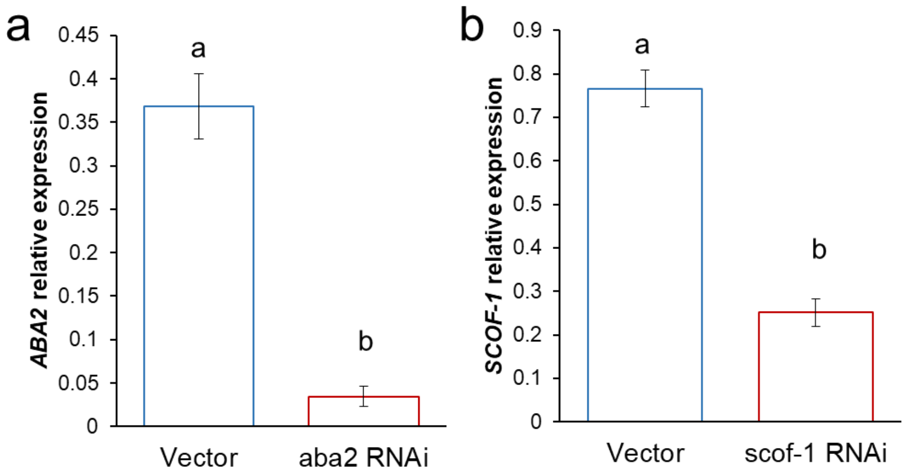


**Fig. S2. VIGS knockdown confirmation.** Basal expression levels of ABA2 (**a**) or SCOF-1 (**b**) were assayed via quantitative PCR in plants infected with a vector control or with vectors for *aba2* RNAi or *scof-1* RNAi, respectively. Different letters indicate significance at $P<0.05$, LSD multiple comparisons test.

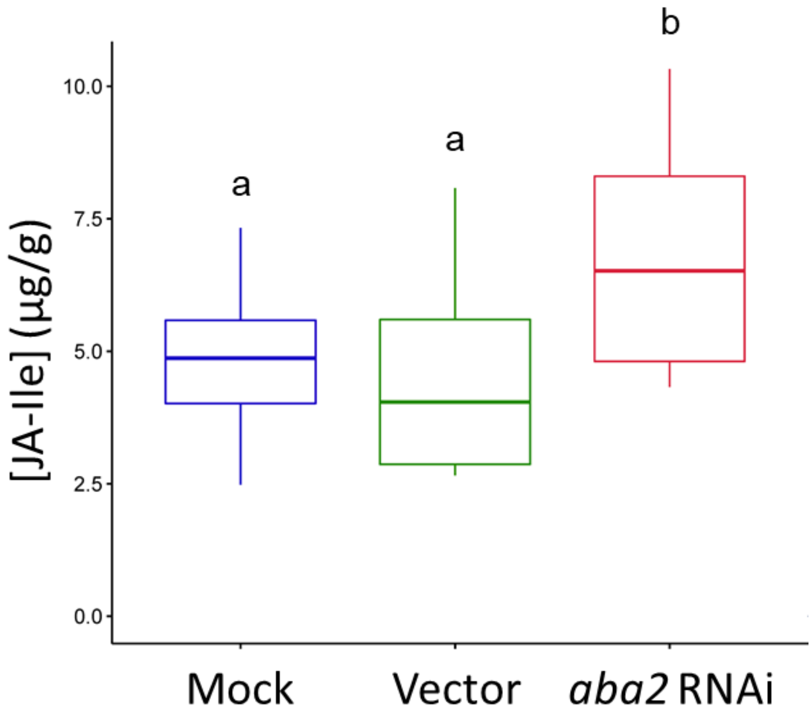


**Fig. S3. Accumulation of JA-Ile in response to aphid feeding in control and ABA-deficient plants.** Leaf samples were collected from plants infested with aphid for seven days. After aphid removal, leaves were extracted and JA-Ile was quantified by liquid chromatography– mass spectrometry (LC-MS). Different letters indicate significance at $P<0.05$, LSD multiple comparisons test. Data from two independent experiments were included in the analysis.

**Table S1. Primers used to generate RNAi constructs and quantify gene expression (qRT-PCR).**

| Gene | Glyma ID | F primer 5'-3' | R primer 5'-3' |
|---|---|---|---|
| **Gene Silencing (RNAi) constructs** | | | |
| *aba2*-RNAi | Glyma.11g151400 | AAATGGATCCAAGGAAGAGCACAGCATTAGC | ATATCTCGAGAAGAAGGGCTCAATCATTTCTTT |
| *scof-1*-RNAi | Glyma.17g236200 | AAATGGATCCGAGTTTCCGGTGACTGGCC | ATATCTCGAGCCATCTTTTCCCTTTGACGACC |
| **Quantitative RT-PCR** | | | |
| *ABA2* | Glyma.11g151400 | CTCCAACACAAAGGCTATTAGG | CATTGTTGACTATGATGTGAAGG |
| *SCOF*-1 | Glyma.17g236200 | CCATCTTTTCCCTTTGACGA | TTTCCGTTACCGTTACCTT |
| *PinN2* | Glyma.14g038300 | CGCTCTAGAGAAAGTGCAAGAATTA | CTCACCCAAACCTTCGCTTC |
| *CYP94C1a* | Glyma.12g087200 | AAGGGTGTGTTTGGGGAAGG | CACAAACCCGAACCGGAAAC |
| *CYP94C1b* | Glyma.11g185700 | CGAAAGATGGCTGCGTGATG | CCTTCCCCAAACACACCCTT |
| *UBQ* | Glyma.20g141600 | TCTCCCTTCAAGATGCAGA | GAGGTGAAGAGTACTCTCCTT |

**Table S2. Average (± SEM) phytohormone content in control and aphid-infested plants 7 days after infestation.** a=ng/g fresh tissue; b=μg/g dry weight; * indicates aphid-infested plants had significantly (P<0.05) altered hormone content compared to control plants

| Phytohormone | Control | Aphid-Infested |
|---|---|---|
| *cis*-JA[a] | 3.55 ± 1.45 | 19.82 ± 4.72* |
| *trans*-JA[a] | 68.45 ± 6.49 | 70.54 ± 6.73 |
| JA-Ile[b] | 4.53 ± 0.43 | 4.83 ± 0.52 |
| ABA[b] | 5.34 ± 0.36 | 7.67 ± 0.77* |
